# An *In silico* Algorithm for Identifying Amino Acids that Stabilize Oligomeric Membrane-Toxin Pores through Electrostatic Interactions

**DOI:** 10.1101/716969

**Authors:** Rajat Desikan, Prabal K. Maiti, K. Ganapathy Ayappa

## Abstract

Pore forming toxins (PFTs) are a class of proteins which have specifically evolved to form unregulated pores in target plasma membranes, and represent the single largest class of bacterial virulence factors. With increasingly prevalent antibiotic-resistant bacterial strains, next generation therapies are being developed to target bacterial PFTs rather than the pathogens themselves. However, structure-based design of inhibitors that could block pore formation are hampered by a paucity of structural information about pore intermediates. On similar lines, observations of the inter-subunit interfaces in fully-formed pore complexes to identify druggable residues, whose interactions could potentially be blocked to hamper pore formation or destabilize pore assemblies, are often limited because of the presence of a large number of protein-protein interaction sites across pore inter-subunit interfaces. Narrowing down the list of plausible target residues requires a quantitative assessment of their contributions towards pore stability, which cannot be gleaned from a single, static, crystal or cryo-EM pore structure. We overcome this limitation by developing an *in silico* screening algorithm that employs fully atomistic molecular dynamics simulations coupled with knowledge-based screening to identify residues engaged in persistent and stabilizing electrostatic interactions across inter-subunit interfaces in membrane-inserted PFT pores. Application of this algorithm to prototypical *α*-PFT (cytolysin A) and *β*-PFT (*α*-hemolysin) pores yielded a small predicted subset of highly interacting residues, blocking of which could destabilize pore complexes as shown in previous mutagenesis experiments for some of these predicted residues. The algorithm also yielded a novel set of residues in both cytolysin A and *α*-hemolysin pores for which no mutagenesis and stability data exists to the best of our knowledge, and therefore could serve as hitherto un-recognised potential targets for PFT inhibitors. The algorithm worked equally well for both *α* and *β*-PFT pores, and could thus be potentially applicable to all pores with known structures to generate a database of pore-destabilizing mutations, which could then serve as a starting point for experimental validation and structure-based PFT-inhibitor design.

## 1. Introduction

The cellular plasma membrane, which is mainly comprised of a few nanometre thick lamellar sheet of amphipathic lipids, carbohydrates and bound membrane proteins, serves as a semi-permeable barrier that actively regulates the transport of ions and small molecules from the cytosol to the extra cellular environment and vice versa (Phillips et al., 2009). Therefore, a structurally intact plasma membrane, which carefully facilitates and regulates selective transport in response to various signals, is important for the functioning of vital cellular processes and thus survival. To exploit this vulnerability, molecular weapons produced by many organisms in the animal kingdom, from pathogenic bacteria to sea anemones, comprise proteins known as pore forming toxins (PFTs), which specialize in nullifying the ability of target plasma membranes to regulate selective transport (Peraro & van der Goot, 2016; M. W. Parker & Feil, 2005; Los et al., 2013). In fact, PFTs are the main virulence factors of pathogenic bacteria that cause most human diseases, such as *Vibrio cholerae, Staphylococcus aureaus, Pseudomonas aeruginosa, Bacillus anthracis*, and others. With the increase in the prevalence of human and animal infections propagated by antibiotic-resistant bacterial strains (Blair et al., 2015; Opal, 2016), an important research focus is on targeting bacterial virulence factors rather than the pathogens themselves (Gammon, 2014; Escajadillo & Nizet, 2018). Since PFTs form the single largest class of bacterial virulence factors (Los et al., 2013), PFTs of pathogenic bacteria are currently being targeted by many novel therapies, and therefore, an understanding of the molecular features and mechanisms of PFTs are crucial for such therapies to succeed.

PFTs disrupt the structural integrity of the host cell membrane at the molecular level via the following mechanism. PFTs from the virulent organism are secreted as water soluble ‘monomers’, that conformationally transform (Peraro & van der Goot, 2016; Giri Rao et al., 2016) upon encountering the host membrane into ‘protomers’ and circularly oligomerize to form nanopores, either directly or via a non-membrane-inserted prepore intermediate that later punches into the membrane (Peraro & van der Goot, 2016). Formation of sufficient numbers of these multimeric and mostly non-selective nanopores on the plasma membrane eventually leads to the disruption of vital signalling processes, cripples the selective permeability of the plasma membrane by causing unregulated transport of ions and small molecules, and causes cell death (Peraro & van der Goot, 2016; M. W. Parker & Feil, 2005), systemic inflammation (Los et al., 2013; Fang et al., 2017) and propagation of infection (Czuczman et al., 2014; Zafar et al., 2017). PFTs are mainly classified on a structural basis as *α*-PFTs or *β*-PFTs based on whether amphipathic *α*-helices or *β*-hairpins form the dominant secondary structure of the pore trans-membrane domain (M. W. Parker & Feil, 2005). A few solved *α* and *β* PFT nanopore structures from various organisms are illustrated in Figure 1. Most PFT pores are a few nanometres in size, with the largest PFT pores ranging between 30 *−* 50 nm in diameter (Peraro & van der Goot, 2016; Los et al., 2013).

**Figure 1.**
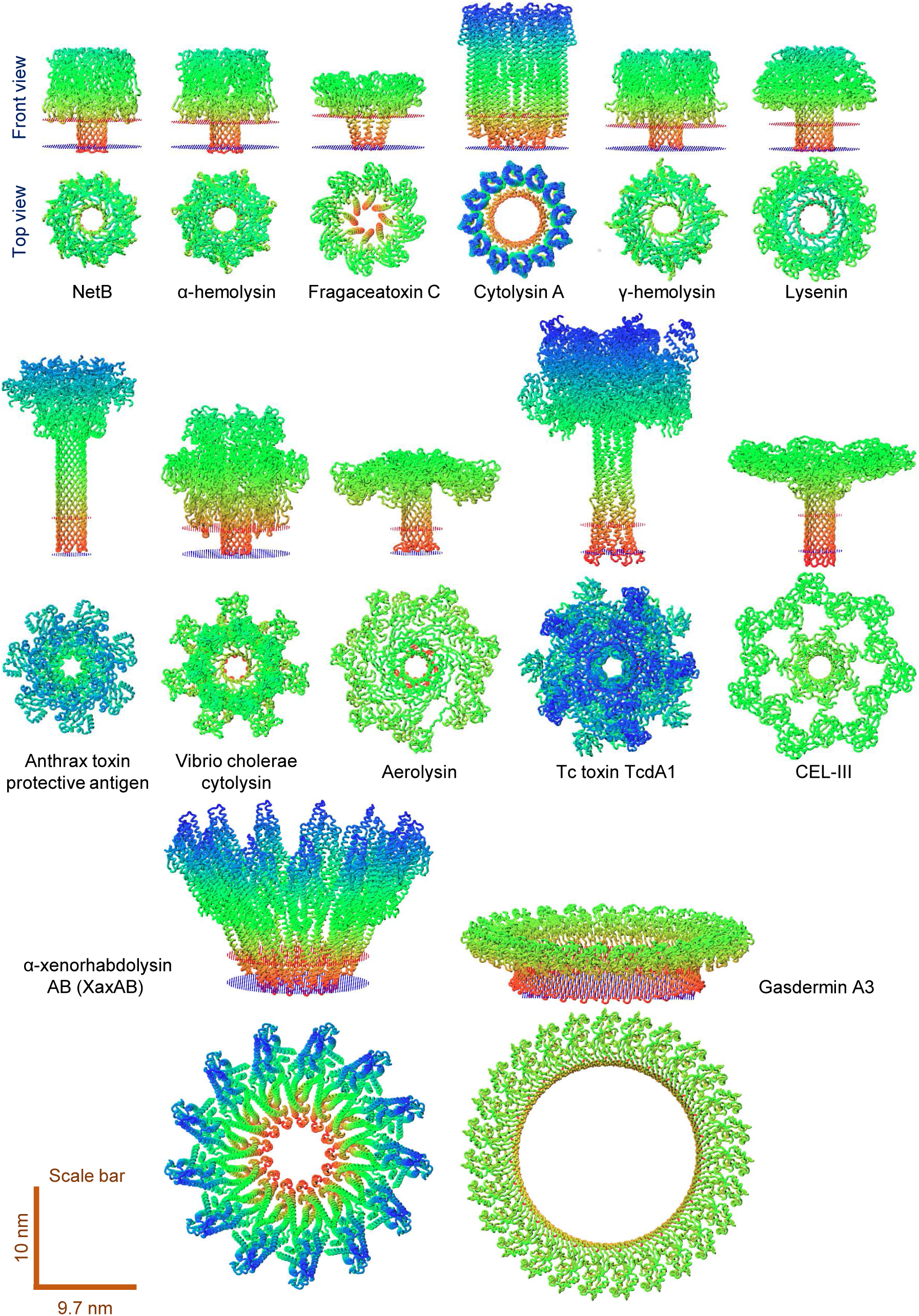
Front and top views of known *α* and *β* PFT structures from various organisms are shown with relative sizes set to scale (scale bars at the bottom). The transmembrane domains are defined from the ‘orientations of proteins in membranes (OPM) database’ (Lomize et al., 2012), and are demarcated by sheets of dummy atoms coloured red and blue for the extracellular-facing and cytoplasm-facing leaflets, respectively. Fragaceatoxin C, cytolysin A and *α*-xenorhabdolysin AB are *α*-PFTs while the rest are *β*-PFTs. Starting from the top left, the PDB IDs are: 4H56, 7AHL, 4TSY, 2WCD, 3B07, 5GAQ, 3J9C, 3O44, 5JZT, 5LKI, 3W9T, 6GY6, and 6CB8.

To target PFTs directly, small molecules, peptides and liposome decoys that inhibit the binding of PFTs to host receptors, oligomerization of the pores on the host plasma membrane, or block already formed pores have been designed (recently reviewed in Escajadillo & Nizet, 2018) to protect against infection propagated by PFTs of *Bacillus anthracis* (anthrax) (Nestorovich & Bezrukov, 2014; Rai et al., 2006), *Staphylococcus aureaus* (pneumonia, gut and skin infections) (Ragle et al., 2010), *Escherichia coli* (gut and urinary tract infections) (Mandal et al., 2016), *Vibrio cholerae* (cholera) (Rai et al., 2006), *Streptococcus pneumoniae* (pneumonia) (Henry et al., 2015), and many other bacteria. While inhibitors that block fully formed pores may yield some therapeutic benefit, blocked pores can nonetheless cause other disruptive effects such as bending and deformation of the plasma membrane (Tzokov et al., 2006; Drücker et al., 2019) and alteration of the lateral organization of lipid domains (Yilmaz & Kobayashi, 2015) as well as lipid dynamics (Ponmalar et al., 2019). Further, we have also shown through experiments and modelling that not just fully formed pores but oligomeric intermediates along the pore formation pathway of the *α*-PFT, Cytolysin A, are also capable of spontaneously compromising membrane integrity by causing leakage (Agrawal et al., 2017; Desikan, Maiti, & Ayappa, 2017). Therefore, rather than blocking fully formed pores, a more effective strategy to neutralize PFTs would be to prevent pore formation, either by shielding the host plasma membrane from PFTs, or by blocking oligomerization and thus pore formation. This strategy underlies the design of therapeutics such as toxin-absorbing nanosponges, which act as a sink for PFTs and therefore protect the host cell membrane (Henry et al., 2015; Hu et al., 2013), small molecule inhibitors of PFT oligomerization (Escajadillo & Nizet, 2018), passive immunization with PFT-targeting antibodies (Foletti et al., 2013; Tkaczyk et al., 2016), PFT-receptor blockade on host cell membrane that prevents membrane-binding and conformational change (Rosch et al., 2010; Johnson et al., 2013), and even vaccination strategies with PFT epitopes as antigen that could promote anti-PFT adaptive-immune responses preventing the binding and oligomerization of PFTs (Adhikari et al., 2016; Wei et al., 2017).

Effective inhibitors that target PFT oligomerization and pore formation should bind to critical residues on membrane-bound PFT protomers and block protomer-protomer interfacial interactions, thus impairing self-assembly and the stability of these oligomeric and non-covalently bound protein complexes (Escajadillo & Nizet, 2018). However, identification of these critical residues and their interactions in the inter-protomer interface has been a challenge. Direct visualization of the pore-forming process in molecular detail, especially the membrane-bound protomer and pore-intermediate conformations, is a formidable issue since pores lie within or at the edge of the optical resolution limit. Additionally, temporal resolution of pore formation is tricky since it is a rapid process that typically finishes within time scales of seconds (Thompson et al., 2011; Sathyanarayana et al., 2018). For these reasons, experimentally solved structures of pore intermediates are not available, which precludes direct structure-based inhibitor design against protomers and pore-intermediates. One solution is to solve the inverse problem – screen the fully oligomerized PFT pore structures (Figure 1) for inter-protomer interacting residues that are crucial for pore stability. Designed PFT inhibitors could then target these residues, thus undermining the structural stability of pore-intermediates as well as fully formed pores.

Here, we postulate that buried inter-protomer electrostatic interactions (hydrogen bonds and salt bridges) are crucial for the stability of large, multimeric pore complexes. The support for this argument was based on two observations. Firstly, previous studies have shown that electrostatic interactions such as hydrogen bonds and salt bridges contribute significantly to protein stability and conformational specificity (Desiraju & Steiner, 2001; Waldburger et al., 1995; Bosshard et al., 2004; S. Kumar & Nussinov, 2002; Donald et al., 2011; Basu & Mukharjee, 2017). Secondly, from a representative dataset of 122 non-PFT oligomeric membrane proteins (Gupta et al., 2017), the average number of inter-subunit salt bridges were 3.2 *±* 3.6, whereas representative PFT pores such as *α*-hemolysin, cytolysin A and Tc toxin TcdA1 (Figure 1) engage in 11, 13 and 29 inter-protomer salt bridges, respectively (calculated from the PDBePISA server (Krissinel & Henrick, 2007) with PDB IDs 7AHL, 2WCD and 5LKI). These observed enhanced electrostatic networks between protomers in PFTs, particularly interactions that may occur in the low-dielectric non-solvated protein interface, could be stronger and more specific than solvent exposed electrostatic interactions (Waldburger et al., 1995; Bosshard et al., 2004; S. Kumar & Nussinov, 2002; Donald et al., 2011; Basu & Mukharjee, 2017), and may play key interface-stabilizing roles in facilitating oligomerization, prepore-to-pore transition and pore stability (Wade et al., 2015; Mueller et al., 2009). Additionally, many of the larger PFT pores possess much higher buried protomer-protomer interfacial surface areas than non-PFT membrane proteins – for example, 6570.7 Å^2^ between protomers in the Tc toxin TcdA1 (shown in Figure 1), compared to an average of 2285.1 *±* 1364.0 Å^2^ for non-PFT membrane proteins from the aforementioned dataset. Therefore, we reasoned that buried electrostatic interactions across large PFT interfaces could be instrumental in facilitating specific protomer binding and oligomerization during pore formation as well as the structural stability of fully formed PFT pore complexes.

However, many challenges persist. While proteins are flexible structures that typically exist in an ensemble of conformations, experimentally determined pore structures (Figure 1) are just a single low-energy static conformation that may not be representative of the conformational ensemble. Additionally, the few *α*-PFT and *β*-PFT pore structures which have been elucidated (Figure 1) were not determined in their native membrane environments but in detergents. Detergents have been shown to form dynamic micellar-like structures both in solution and around membrane proteins (Sammalkorpi et al., 2007; Böckmann & Caflisch, 2005), and the choice of detergents used – non-ionic or zwitterionic – may impact protein conformation (Rouse & Sansom, 2015). Hence, observing electrostatic interactions by examining a single detergent-determined PFT pore structure may only yield partial insights; an issue that could be ameliorated by analysing an equilibrium ensemble of membrane-inserted pore conformations. Secondly, PFT pores are large protein complexes with high buried interfacial area, and often, a large number of residues engage in numerous inter-protomer interactions. For example, one of the smaller pores, *α*-hemolysin (Figure 1), has an average of 85 interacting residues in the inter-protomer interface, while a larger pore complex, Tc toxin TcdA1 (Figure 1), has an average of 186 interfacial residues (calculated from the PDBePISA server (Krissinel & Henrick, 2007) with PDB IDs 7AHL and 5LKI). Without information on protein dynamics, it is not possible to accurately assess, rank and screen the contributions of a particular interacting residue towards stabilizing the inter-protomer interface (Donald et al., 2011). Therefore, additional resource and time-consuming experiments such as mutagenesis (alanine/cysteine scanning) (Walker & Bayley, 1995a; Panchal & Bayley, 1995) or advanced mass spectrometry techniques optimized for oligomeric membrane proteins (Gupta et al., 2017) have to be performed for obtaining this information.

In this article, we formulated an *in silico* algorithm that addresses the above technical challenges in order to robustly identify critical interfacial residues involved in pore-stabilizing inter-protomer electrostatic interactions. In the following sections, we describe the principles underlying the *in silico* algorithm, employ the proposed algorithm on representative *α*-PFT and *β*-PFT pores to predict critical stabilizing residues, and compare the predictions with previously reported experimental data for validation.

## 2. Results

### 2.1. Computational algorithm for identifying interface-stabilizing salt bridges and hydrogen bonds

We constructed a stepwise *in silico* algorithm (Figure 2) with the aim of robustly identifying residues that stabilized the inter-protomer interface in pores via buried electrostatic interactions such as hydrogen bonds and salt bridges. We tested this algorithm on two toxin pores: (i) the dodecameric *α*-PFT *Escherichia coli* Cytolysin A (ClyA) pore, and (ii) the heptameric *β*-PFT *Staphylococcus aureus α*-hemolysin (AHL) pore (Figures 1 and 2). We choose these as representative pores from the *α*-PFT and *β*-PFT classes, having different structural topographies (Figure 1). A test of robustness of the algorithm is to accurately identify interface-stabilizing residues in pores from both classes of PFTs. The algorithm is described below and schematically illustrated in Figure 2.

**Figure 2.**
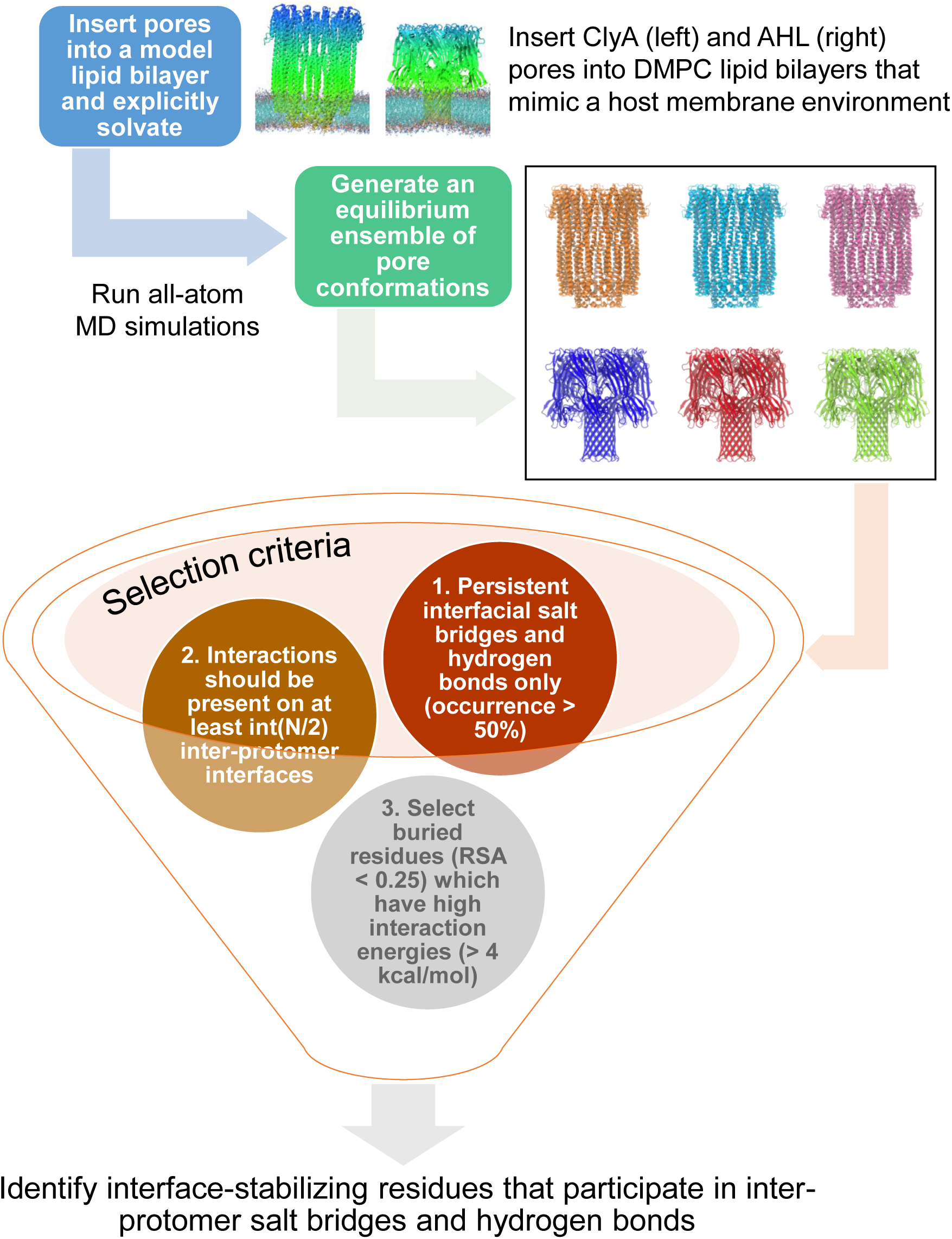
The computational screening algorithm for identifying important interfacial electrostatic interactions in ClyA and AHL pores (see text for details).

The detergent-stabilized ClyA and AHL pore structures are initially inserted into model 1,2-dimyristoyl-sn-glycero-3-phosphocholine (DMPC) lipid bilayers, that physicochemically resemble the host plasma cell membrane environment. To obtain a converged structural ensemble of pore conformations with atomistic insight into protein dynamics, we performed unbiased all-atom molecular dynamics (MD) simulations of the membrane-inserted pores in explicit solvent at physiologically relevant conditions, and analysed protein conformations for hydrogen bonds and salt bridges. Hydrogen bonds were considered to exist if the donor-acceptor distance is less than a cut-off of 3 Å and the hydrogen-donor-acceptor angle is less than a cut-off of 20*°*. Salt bridges were considered to exist if any oxygen atom of an acidic residue (aspartic acid, glutamic acid) fell within a cut-off of 4 Å (Barlow & Thornton, 1983) from any nitrogen of a basic residue (lysine, arginine, histidine). From analysing the MD generated pore ensembles with these definitions, a large number of unique hydrogen bonds and salt bridges that occurred in any pore conformation were catalogued and subjected to the screening funnel (Figure 2).

The screening funnel applied a three-step elimination procedure for the above list of electrostatic interactions. Firstly, only persistent and not transient interactions were chosen. Persistence was quantified from the pore ensembles by employing occurrence frequencies (see ‘Materials and Methods’). Any interaction that had an occurrence of less than a chosen threshold of 50% were eliminated. Additionally, since only interface-stabilizing interactions that ensured the structural integrity of the multimeric pores were of interest, intra-protomer interactions which could play an indirect role in interfacial stability by stabilizing the individual protomer conformations were discarded. Secondly, the high frequency hydrogen bonds and salt bridges from the previous step were checked for their global effects on the pore structure. Since pores are mostly homo-oligomers (see Figure 1; bi-component toxins such as *γ*-hemolysin and *α*-xenorhabdolysin are exceptions), each protomer-protomer interface in the pore complexes is repeated *N* times, where *N* is the number of protomers comprising the pore (for example, *N* = 12 for ClyA and 7 for AHL). Therefore, each inter-protomeric hydrogen bond and salt bridge should in principle be symmetrically repeated multiple times across the interfaces. However, with weakly interacting residues, pore dynamics and breathing motions may lead to fluctuating hydrogen bonds and/or salt bridges. Thus, it is possible that local persistent interactions may occur in only a few inter-protomer interfaces, and these interactions may not contribute across protomers towards the global structural stability of the pore complexes. To unambiguously select highly stabilizing residues, we apply the following screening criteria: any interaction between residues across an inter-protomer interface have to be present in at least 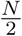 pore interfaces if *N* is even, or 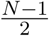 interfaces if *N* is odd. Thirdly, as discussed in the introduction, only hydrogen bonding and salt bridging residues that were buried in the solvent-inaccessible portion of the inter-protomer interfaces, i.e., relative solvent accessibility of that residue (*RSA*) was lesser than 0.25, and had high interfacial interaction energies as computed from the MD generated ensemble (*< −*4 kcal/mol; see Results subsection titled ‘Screening selected amino acid residues using interaction energies and relative solvent accessibilities’), were identified as interface-stabilizing residues.

### 2.2. Molecular dynamics simulations of ClyA and AHL pores in a DMPC membrane

To generate an ensemble of equilibrated pore conformations and to track protein dynamics in membrane environments, we performed 500 ns long atomistic MD simulations of the ClyA and AHL pores in a DMPC lipid bilayer. Analysis of measures that quantified global structural integrity enabled identifying the portion of MD trajectories where simulated pore conformations were fully equilibrated with their environment. Figure 3*a, b* show time traces of backbone root mean square deviation (RMSD) of both pores from their initial structures at 0 ns, and the radius of gyration (*R*_*g*_). It can be observed that both quantities start plateauing at 450 ns, although the equilibrium RMSD of ClyA (3.68 *±* 0.06 Å) was higher than that of AHL (1.45 *±* 0.04 Å), implying that the AHL pore was structurally more stable compared to the ClyA pore. Also shown in Figure 3*b* are the superimposed conformations of single protomer chains from the ClyA and AHL pores, where blue depicts the crystal structure, and red, the final MD snapshot at 500 ns, respectively. It can be observed that the backbone structural deviations are minor, and most changes arise from disordered loops present at the termini and at various junctions connecting *α*-helices and *β*-sheets. Therefore, pore conformations between 450 *–* 500 ns (lilac shaded region in Figure 3*a, b*) were extracted to construct the equilibrium pore conformational ensembles for analysing hydrogen bonds and salt bridges.

**Figure 3.**
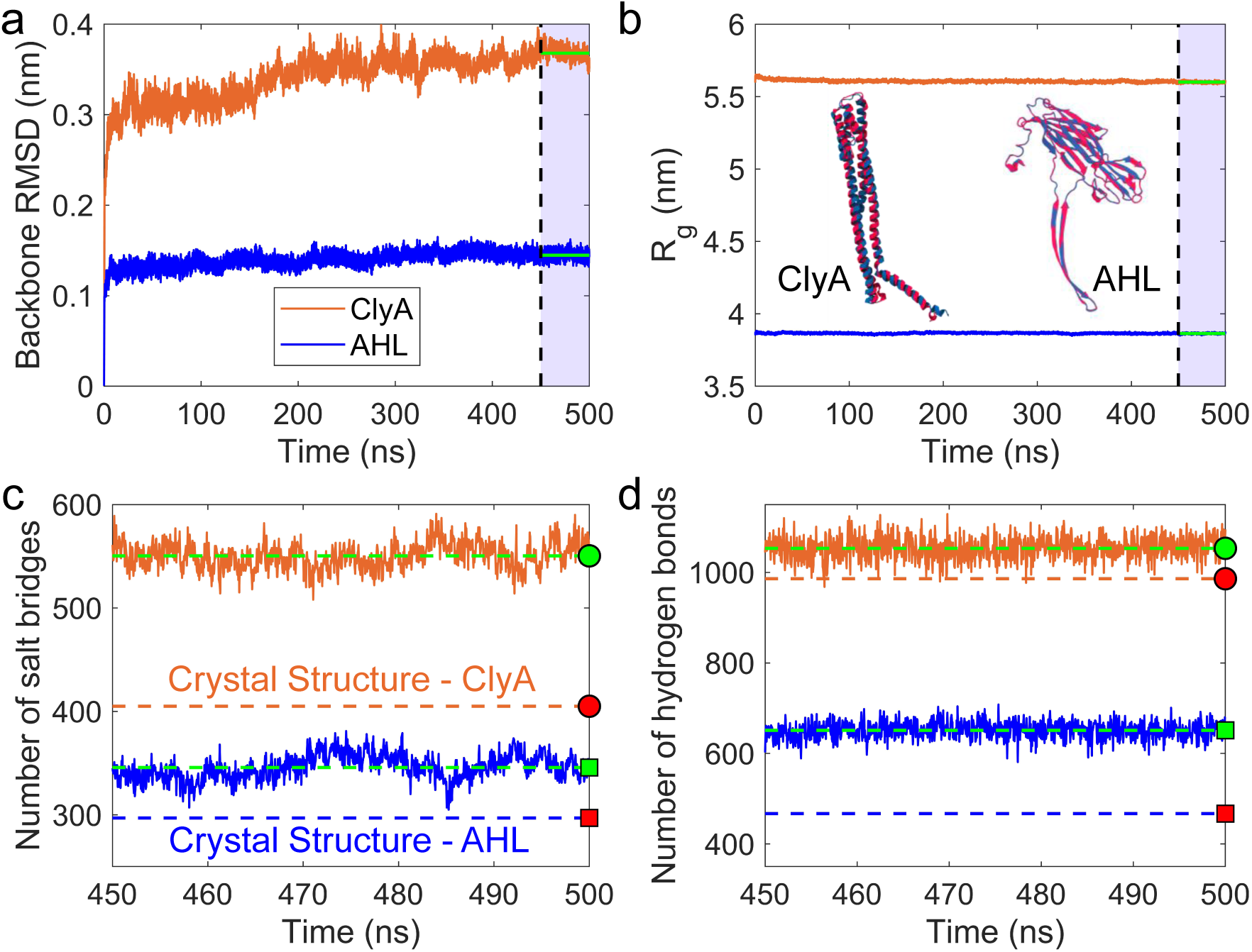
MD simulations of ClyA and AHL pores in a DMPC membrane. (**a**) Time traces of backbone root mean square deviation (RMSD) of both pores from their initial structures at 0 ns. The lilac highlighted region from 450-500 ns indicates the plateau where the pore conformations have relaxed (mean RMSD in this region shown as a thick green line), and this portion of the trajectories yielded the equilibrium pore conformational ensembles for further analysis. (**b**) Time traces of radius of gyration (*R*_*g*_). Also shown are superimposed protomer conformations from the respective pore crystal structures (blue) and the final MD snapshots at 500 ns (red). Time trends of the overall (including both intra and inter protomer) number of (**c**) salt bridges and (**d**) hydrogen bonds from the equilibrated portion of the trajectories. MD averages were marked by green dashed lines and indicated on the y-axis by green filled circles and squares for ClyA and AHL, respectively. The number of salt bridges and hydrogen bonds in the pore crystal structures are shown by orange (ClyA) and blue dashed lines (AHL), and indicated on the y-axis by either red filled circles (ClyA) or squares (AHL).

Analysis of local side-chain rearrangements revealed that compared to their crystal structures, both ClyA and AHL showed a significant increase in the overall number of salt-bridges (Figure 3*c*). The crystal structure of the ClyA pore exhibited 405 salt bridges, while the MD generated ensemble had an average of 550 *±* 13 salt bridges, indicating a 36% increase. Similarly, the crystal structure of the AHL pore exhibited 297 salt bridges while the MD generated ensemble had an average of 346 *±* 12 salt bridges, indicating a 17% increase. Similar observations have been made for other systems – salt bridges which were previously not present in the crystal structures of proteins were observed to be formed during the course of an MD simulation (Karshikoff & Jelesarov, 2008). Figure 3*d* shows that the number of intra-protein hydrogen bonds in the AHL crystal structure and the MD generated ensemble were 986 and 1053 *±* 25 respectively, and that in the AHL crystal structure and the MD generated ensemble were 467 and 651 *±* 19 respectively, indicating a 6% and 39% increase for the MD simulated conformations of ClyA and AHL, respectively. These observations point towards the crucial role of membrane-solvent interactions in enhancing the stability of membrane-inserted pore conformations compared to their crystal structures determined in detergents.

### 2.3. Identifying persistent interfacial salt bridges and hydrogen bonds from occurrence frequencies

From the equilibrium MD generated ensemble of pore conformations, we catalogued all unique hydrogen bonds and salt bridges that occurred in the pores and began selecting interactions according to the screening funnel shown in Figure 2. We recorded a total of 11912 and 6438 hydrogen bonds, 1776 and 1039 salt bridges, between unique sets of interacting atoms in the ClyA and AHL pores, respectively. As per the first screening criteria, we wished to identify persistent and not transient interactions. To quantify persistence, we computed occurrence frequencies of hydrogen bonds and salt bridges from the MD conformations. The occurrence frequency (*ν*) of a particular hydrogen bond or salt bridge was defined as the fraction of pore conformations in the equilibrium ensemble in which that particular interaction was switched on:

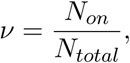

where *N*_*on*_ is the number of frames in which the salt bridge is formed, and *N*_*total*_ = 1500 is the total number of frames in the ensemble. The concept of occurrence is illustrated in Figure 4 by considering three sets of D-K salt bridges in ClyA as examples. As shown in Figure 4*a*, D has two negatively charged oxygen atoms that could both form salt bridges with the positively charged nitrogen atom in K. The distances between the nitrogen atom in K and the two oxygen atoms in D were denoted as Δ_1_ and Δ_2_. For the inter-protomer salt bridge in ClyA, D74-K240, both Δ_1_ and Δ_2_ were less than the salt bridging threshold of 4 Å (Barlow & Thornton, 1983) in all the equilibrated pore conformations, thus leading to 100% occurrence. In contrast, for the intra-protomer salt bridges D25-K29 and D7-K14, Δ_1_ and Δ_2_ frequently rose above the 4 Å threshold thus resulting in the salt bridge interactions switching between ‘on’ and ‘off’ states (Figure 4*c, d*). This resulted in occurrence frequencies of 42.9 % and 5.4 % for D25-K29 and D7-K14 respectively. Thus, D25-K29 and D7-K14 formed transient salt bridges whereas D74-K240 formed a persistent salt bridge. These observations emphasize the advantages of analysing protein dynamics from a conformational ensemble to track ‘on-off’ switching events and subsequently distinguish between transient and persistent electrostatic interactions rather than simply observing static non-covalent bonds in crystal/cryo-EM structures.

After occurrence frequencies were computed for all unique hydrogen bonds and salt bridges, they were binned into histograms as shown in Figure 5*a-d*. A majority of the interactions were transient and possessed occurrence frequencies *<* 50%. Only persistent hydrogen bonds and salt bridges with occurrence frequencies *>* 50% (green shaded regions in Figure 5) were selected for subsequent screening. An analysis of cumulative densities (red dot-dash lines in Figure 5*a-d*) showed that in ClyA, only 13.8% hydrogen bonds and 19.3% salt bridge interactions were persistent, while in AHL, 12.2% hydrogen bonds and 20.0% salt bridges were persistent. These persistent electrostatic interactions were listed, and the list was further pruned by employing two criteria serially: (i) only inter-protomer hydrogen bonds and salt bridges were selected, and (ii) as per screening criteria 2, interactions that did not occur in at least 6 and 3 interfaces of the homo-oligomeric ClyA and AHL pores were discarded.

**Figure 4.**
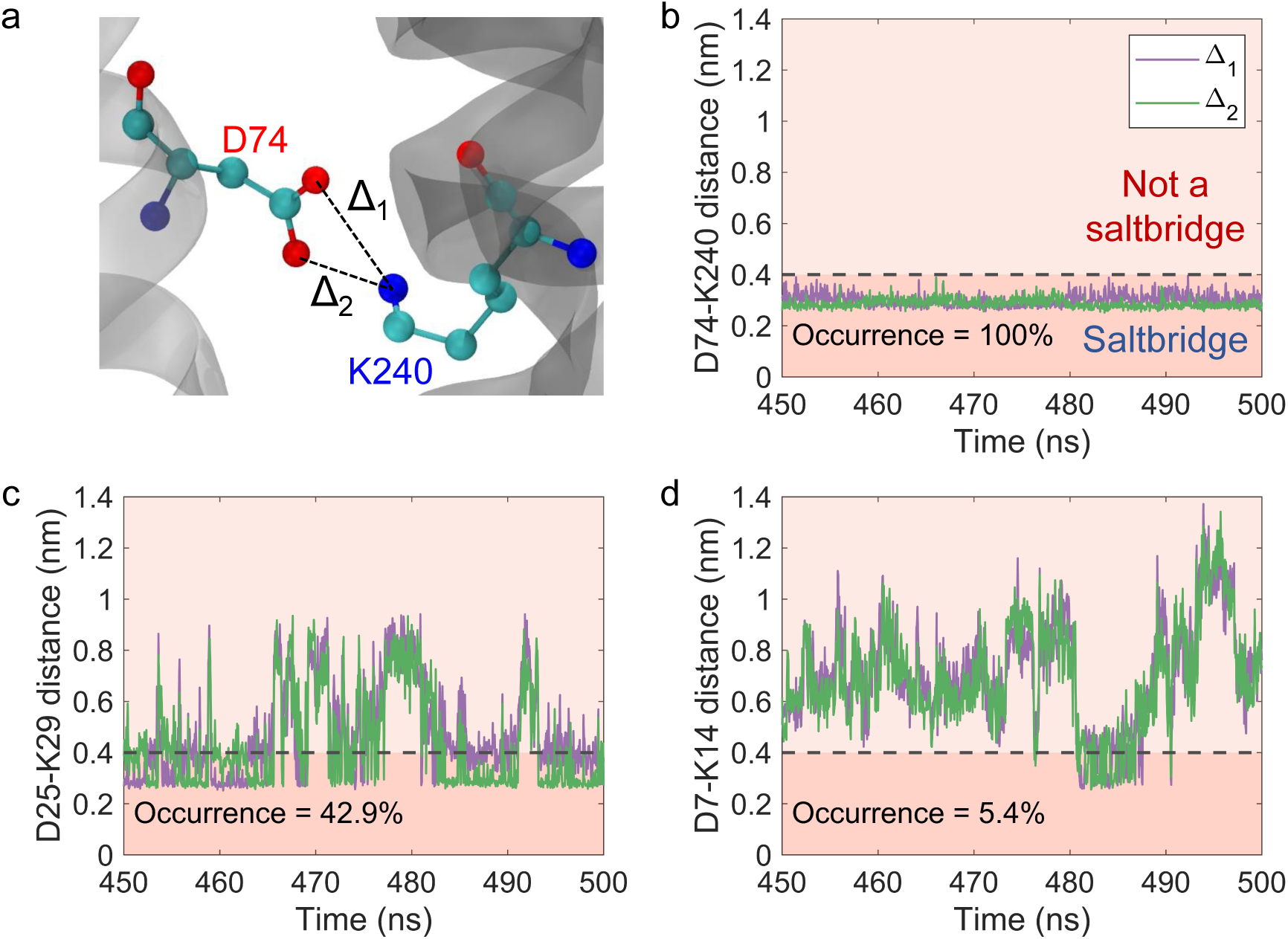
Salt bridges with different occurrence frequencies switching on and off. (**a**) Two inter-protomer salt bridges between D74 and K240 in ClyA are shown. Colour codes: 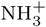 in blue, O^*−*^ in red and carbons in cyan. Hydrogen atoms are not shown for clarity. The oxygen-nitrogen distances were denoted as Δ_1_ and Δ_2_. (**b-d**) To illustrate the concept of occurrence frequencies and salt bridges switching on and off, time traces of Δ_1_ (purple) and Δ_2_ (green) for three D-K salt bridges in the ClyA pore with different occurrences – D74-K240 (100% occurrence), D25-K29 (42.9% occurrence) and D7-K14 (5.4% occurrence) – are shown. The salt bridge ‘on’ (Δ*≤* 0.4 nm; dark orange) and ‘off’ (Δ *>* 0.4 nm; light orange) states demarcated by the 4 Å threshold (horizontal black dashed line) are highlighted.

**Figure 5.**
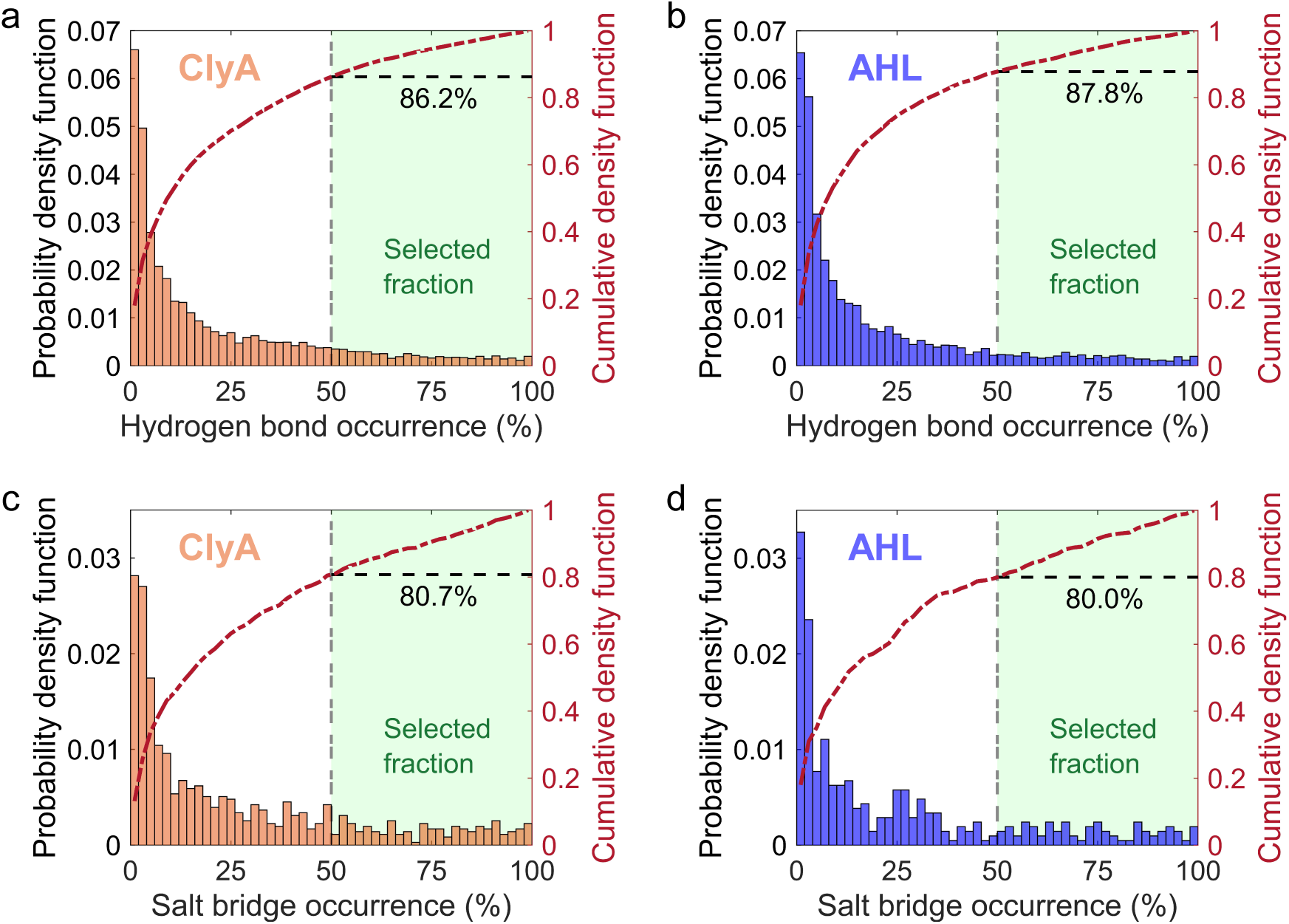
Occurrence frequency distributions for electrostatic interactions in ClyA and AHL pores. (**a, b**) Distribution of hydrogen bond occurrence frequencies, and (**c, d**) distribution of salt bridge occurrence frequencies in ClyA and AHL pores, respectively. Only interactions with occurrence frequencies *>* 50% (green shaded region, threshold marked by a vertical grey dashed line) were selected. Normalized cumulative densities are shown as red dot-dash lines, and the cumulative densities at 50% occurrence are marked by horizontal black dashed lines. The numbers underneath these black dashed lines represent the fraction of transient hydrogen bonds and salt bridges in ClyA and AHL.

### 2.4. Screening selected amino acid residues using interaction energies and relative solvent accessibilities

Application of screening criteria 1 and 2 (see Figure 2) led to a small subset of persistent, inter-protomer, and prevalent hydrogen bonds and salt bridges, which were then subjected to screening criteria 3. As explained in previous sections, criteria 3 only selected buried and strongly interacting residues with non-bonded energies *< −*4 kcal/mol. To classify residues as buried, the per-residue solvent accessible surface area (SASA) was computed for these residues in each protomer of both pores, and the mean SASA for these residues were normalized with the theoretically estimated maximum per-residue SASA (Tien et al., 2013) to obtain the per-residue relative solvent accessibility (RSA). We then employed an oft-used RSA based binary classification of residues as buried and solvated – residues with RSA*≤* 0.25 (or 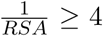) were considered as buried while the rest were assumed to be solvated (Levy, 2010). Similarly, we computed the per-residue total non-bonded interaction energy between each interface in both pores, taking into account contributions from both Coulomb’s and van der Waal’s interactions. For the *i*^*th*^ pore residue with *n* atoms interacting across the *j*^*th*^ inter-protomer interface, the per residue energy, 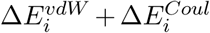, which is averaged across all the inter-protomer interfaces of the pore, is computed using

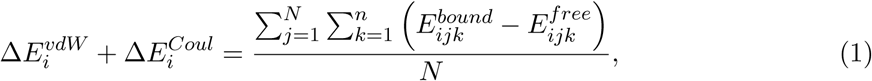

where *N* is the oligomeric state of the pore complex. 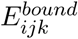 and 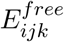 are the non-bonded energies of the *k*^*th*^ atom from the *i*^*th*^ residue at the *j*^*th*^ inter-protomer interface, when the interacting protomers exist as complexed (bound) and free (unbound) states, respectively. We then computed the mean of Δ*E*_*i,j*_ across all protomer-protomer interfaces in both pores to arrive at the per-residue contribution to the interfacial binding energy 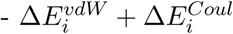 (shown in figures without the subscript *i* for ease of notation).

After computing the energetics for all residues, a ternary classification was made: (i) for a particular residue, if Δ*E*^*vdW*^ +Δ*E*^*Coul*^ *>* 0 kcal/mol, that residue was labelled as ‘destabilizing’, (ii) if *−*4 *<* Δ*E*^*vdW*^ +Δ*E*^*Coul*^ *<* 0 kcal/mol, that particular residue was termed as ‘weak’ because its energetic contribution was less than the bond energy of a weak hydrogen bond (Desiraju & Steiner, 2001), and (iii) if Δ*E*^*vdW*^ + Δ*E*^*Coul*^ *< −*4 kcal/mol, that particular residue was considered to significantly contribute to interface stabilization.

As shown in Figure 6*a, b*, only buried and strongly interacting residues (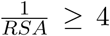 and Δ*E*^*vdW*^ + Δ*E*^*Coul*^ *< −*4 kcal/mol; green shaded region) were identified as interface-stabilizing residues in the ClyA and AHL pores, respectively. These critical residues along with their inter protomer hydrogen bonds and salt bridges at the protomer-protomer interfaces are illustrated in Figure 6*c, d* for ClyA and AHL, respectively. Interestingly in ClyA, some of the salt bridge networks include both stabilizing and destabilizing residues, such as D41-K172-D171-K206, E85-Q131-K254, and E161-K214 (Figure 6*a, c*). However, the net interaction energy for all three networks, calculated by summing the per-residue energies of all interacting amino acids, was favourable and observed to be *−*16.0, *−*11.2, and *−*18.4 kcal/mol for D41-K172-D171-K206, E85-Q131-K254, and E161-K214, respectively.

**Figure 6.**
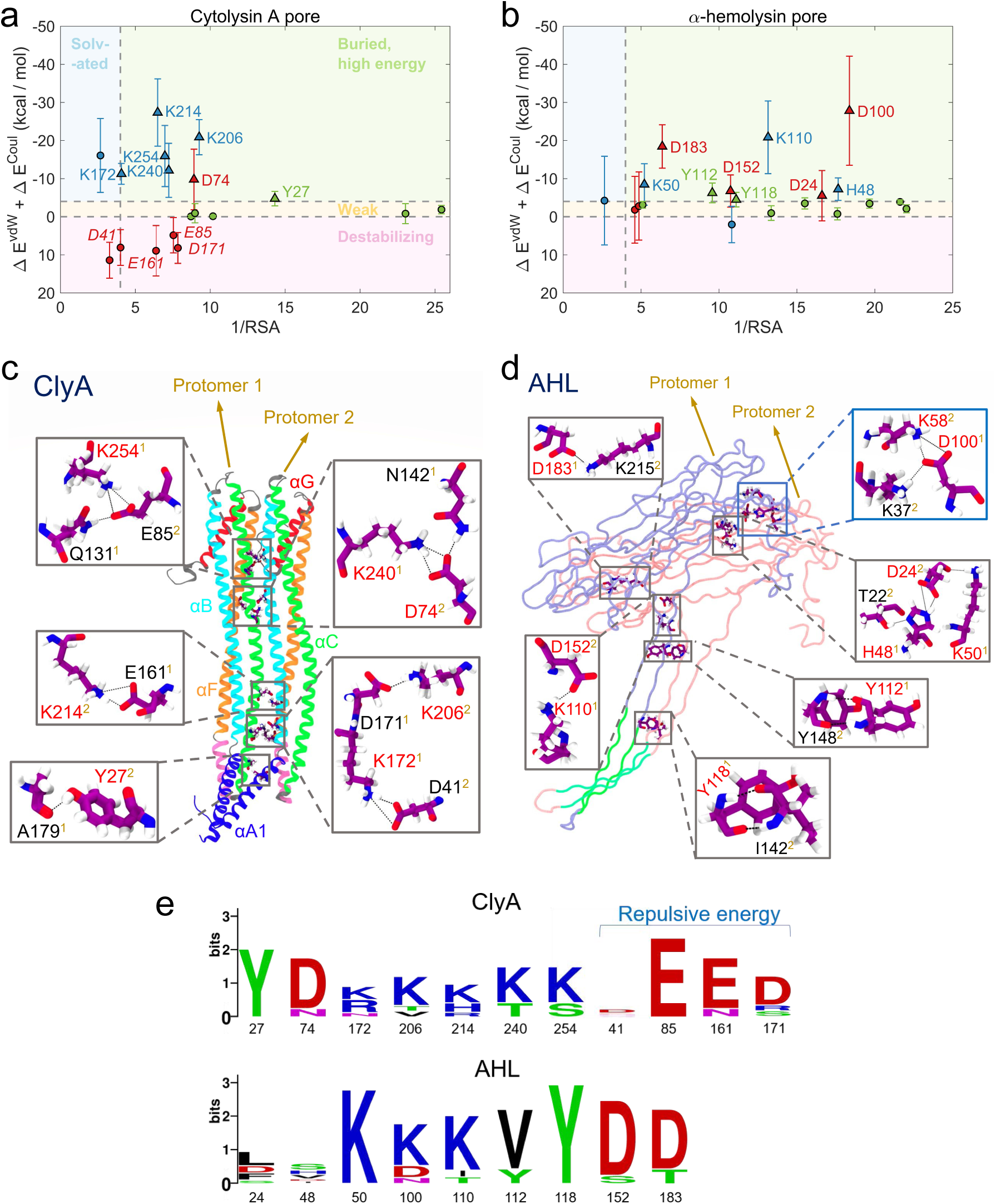
Selection of buried residues that strongly contribute to the interfacial energy in (**a**) ClyA and (**b**) AHL pores based on relative solvent accessibility (RSA) and the per-residue non-bonded interaction energy (Δ*E*^*vdW*^ + Δ*E*^*Coul*^) at the interface. 25% RSA threshold (vertical grey line) and the energy thresholds separating strong (*< −*4 kcal/mol; buried and solvated shown as green and blue shaded regions respectively), weak between (0 and *< −*4 kcal/mol; yellow shaded region) and destabilizing (*>* 0 kcal/mol;pink shaded region, italicized) interactions are illustrated. Only buried residues with high energies (triangles in the green shaded region) are selected and labelled. Colour convention: red circles – negatively charged residues, blue – positively charged residues, and green – others. From the (**c**) ClyA and (**d**) AHL pores, selected residues which satisfy screening criterion 1*−*3 (text coloured red), and their partners which form interfacial electrostatic interactions that satisfy criterion 1 *−* 2 (text coloured black) are shown in two protomers (helices shown with cartoon representation and coloured according to Mueller et al., 2009). AHL is shown with the tube representation. Transmembrane *β*-strands of the two protomers (residues 119 *−* 126, 132 *−* 140) are coloured green, and the other domains are coloured lilac and pink respectively. (one of the overlapping boxes in AHL is coloured blue for contrast.) (**e**) Amino acid frequencies across 7 and 10 aligned homologous sequences for ClyA and AHL are represented as protein sequence logos.

The energy-based classification of residues emerging from the analysis also revealed some distinct differences between ClyA and AHL pores. In the case of ClyA, we observed that about 50% of the final selected residues contributed to a destabilizing effect, while in the case of AHL, although several residues were found to interact in the ‘weak’ regime, only one residue had marginally destabilizing energetics. This is consistent with the inherent structural stability of *β*-PFT pores compared to *α*-PFT pores (see supporting arguments in ‘Discussion and conclusion’, and evidence from RMSD analysis of MD simulations in Figure 3*a*).

### 2.5. Mutagenesis experiments are consistent with predictions

Previous mutagenesis studies on ClyA and AHL validated our predictions of residues that are critical for pore formation and stability. Our screening protocol illustrates that Y27 present on the membrane inserted *α*-helix of ClyA (*α*A1 in Figure 6*c*) forms a hydrogen bond with A179 present on the membrane-inserted hydrophobic motif, which is in close proximity to the helix *α*A1. Recent work from our group has shown that the mutation Y27A reduced the lytic activity of ClyA by 24-fold (Sathyanarayana et al., 2018) and the mutation Y27F completely abolished lytic activity. Further, Y27 was associated with a putative cholesterol recognition and consensus motif (CRAC) located on the *α* helix of ClyA and the neighbouring residues D25 and K29 were associated with strong cholesterol binding sites (Giri Rao et al., 2016; Sathyanarayana et al., 2018). The second set of residues implicated as potential targets (Figure 6*a, c*) are associated with K206 and D171 located on neighbouring membrane inserted *β* tongues at the membrane-water interface. Interestingly, K206 was found to have strong interactions with cholesterol (Sathyanarayana et al., 2018) located in the pocket formed by adjacent *β* tongues with K206 and D171 present on neighbouring protomers that formed the pore. In addition to the previously implicated role played by cholesterol, this study suggests a complex interplay of interactions between *α* helices that contribute to the emergence of stability of the ClyA pore. The rest of the critical residues which lie predominantly in the extracellular space of ClyA pore are potential targets which have not been tested experimentally to the best of our knowledge.

Similarly, in AHL, the formation of the buried salt bridge D152 – K110 is hypothesized to be the electrostatic switch that triggers the pre-pore to pore transition (Wade et al., 2015). Mutating either of these two residues traps AHL in a pre-pore state and abrogates pore formation as well as lytic activity (Panchal & Bayley, 1995; Walker & Bayley, 1995b; Wade et al., 2015; Walker & Bayley, 1995a). Similarly, the mutation, D24C also trapped AHL in a oligomeric prepore state and abrogated lytic activity (Walker & Bayley, 1995a). Mutations H48L, D100C and D183C disrupted electrostatic networks in the inter-protomer interfaces of AHL (Figure 6*d*) and were found to greatly reduce the lytic activity of AHL, thus suggesting impaired pore formation (Walker & Bayley, 1995a; Menzies & Kernodle, 1994). Other predicted critical residues such as K50, Y112 and Y118, represent novel potential mutagenesis targets that have not been verified experimentally to the best of our knowledge.

To estimate the extent to which the predicted residues from our algorithm were conserved in homologous protein sequences, we searched the UniProt database through ConSurf for homologues of chain A from ClyA and AHL, and found 7 and 10 homologous sequences, respectively. These sequences were then aligned and single residue frequencies are illustrated as protein sequence logos in Figure 6*e* (created using the WebLogo server). Most of the final predicted residues in both ClyA and AHL occurred frequently across homologous sequences. Even for residues where substitutions were seen, many of the substituted amino acids had similar electrostatics and chemical nature. For example, the substitution for residue D74 in ClyA was N, and the substitutions for residue K214 in ClyA were H and R, both positively charged salt bridging residues similar to the original K. Interestingly for ClyA, some of the amino acids that possessed destabilizing energies, such as E85 and E161, were also evolutionary conserved, possibly because the net energetic contributions of the electrostatic interaction networks were favourable.

## 3. Discussion and conclusion

Developing inhibitors against the formation of multimeric membrane-inserted pore complexes, which are final self-assembled conformations of bacterial toxins inserted into host cell membranes, is important for developing next-generation therapeutics against increasingly prevalent strains of antibiotic-resistant pathogenic bacteria. Structure-based drug design of inhibitors which target the protomer-protomer interface of pore intermediates may contribute towards therapies that impair oligomerization and pore formation. However, a lack of structural information about oligomeric pore intermediates, and a large number of interacting residues across inter-protomer interfaces in experimentally determined static pore conformations (illustrated in Figure 1 for many PFTs), represent major obstacles towards developing such therapies. We postulated that persistent, de-solvated and strong electrostatic interactions (hydrogen bonds and salt bridges) between two protomer subunits may be important for their oligomerization and pore stability, and hence were potential targets for inhibitor design. The rationale behind this postulate was our observation that PFTs possessed extensive protein-protein interfaces that facilitated a higher number of inter-protomer electrostatic interactions compared to other membrane proteins. Hence, we reasoned that these interactions could facilitate protomer-protomer oligomerization during pore formation as well as the stability of multimeric pore complexes.

On this basis, we developed an *in silico* algorithm (illustrated in Figure 2) that incorporated the generation of an equilibrated pore conformational ensemble in model lipid membranes (Figures 2 and 3), enabled the identification of persistent inter-protomer hydrogen bonds and salt bridges from subtle changes in protein dynamics (Figures 4 and 5), and classified interacting residues based on: (i) relative solvent accessibility as either buried and solvated, (ii) per-residue non-bonded interacting energy as either destabilizing, weak or strong. From a large number of possible interactions (*∼* 1*−*2*×*10^4^) and interfacial residues (*∼* 1*−*2*×*10^2^), our algorithm helped us separate signal from noise and predicted a small subset of inter-protomer residues – 7 and 10 for ClyA and AHL, respectively (Figure 6) – which could be utilized as potential drug targets in the structure-based design of PFT inhibitors, or as plausible mutagenesis targets for altering pore/oligomer stability. Previous data from single-site mutagenesis experiments on ClyA and AHL pores that were available for a subset of the residues predicted by our algorithm confirmed that these residues were indeed important for pore formation and lytic activity. No mutagenesis data exists for the other predicted residues to the best of our knowledge, and we propose that these residues may be hitherto unrecognised novel targets for pore inhibitors and mutagenesis studies of PFTs.

The potential implications of this study are many. Apart from small molecule inhibitor design that targets predicted residues, our algorithm could be employed as a framework for the rational design of single and multiple amino acid PFT mutants that mimic and compete with wild-type PFTs, and dynamically inhibit pore formation. In these engineered mutant protomers, one of the two binding interfaces which dock on to oligomeric pore intermediates would be similar to wild-type, but the other mutated surface would be incapable of forming key inter-protomer interactions and therefore would be unreceptive to further protomer docking. Mixing of mutant PFTs with wild-type could then effectively impede oligomerization and pore formation, since mutants would bind to membrane-bound wild-type protomers and higher order oligomeric intermediates with their unmutated surface, while their other exposed surface would not be conducive for further oligomerization. Such strategies have already been shown to work replacing inter-protomer salt bridges with alanine in a *Staphylococcus aureus* leukocidin, LukGH, led to mutants that could bind to target cells and subsequently oligomerize with the wild-type toxin, thus inhibiting lytic activity (Badarau et al., 2015). Similarly, other approaches have used mutants that bind and oligomerize with wild-type PFTs but trap the assemblies in non-lytic pre-pore conformations, thus conferring protection in both *in vitro* cell-based assays and *in vivo* infection (D. Parker & Prince, 2016; Reyes-Robles et al., 2016). Our algorithm offers a bottom-up approach for rationally ranking and choosing the right inter-protomer residues for efficiently engineering inhibitory PFT mutants that could be employed as therapeutic agents.

On the other end of the spectrum, technologies that rely on nanopore sequencing (H. Kumar et al., 2011) have commonly employed PFT pores such as ClyA (Soskine et al., 2012) and AHL (Clarke et al., 2009). For characterizing proteins, while ClyA nanopores were advantageous compared to *β*-barrel pores such as AHL because of their larger pore lumen, ClyA pores were also more unstable, necessitating directed evolution to produce mutants that could form more stable nanopores (Soskine et al., 2013). This is consistent with oligomeric *α*-helical protein assemblies such as the ClyA pore being less stable than *β*-PFT pores (Gouaux, 1998; Woolfson et al., 2012), especially because their membrane-inserted *α* helices are significantly stabilized by surrounding lipids and cholesterol (Sathyanarayana et al., 2018; Tanaka et al., 2015; Kristan et al., 2009), whereas the rigid *β*-barrels of *β*-PFTs are mainly stabilized by inter-strand interactions of the membrane inserted *β* barrels (Gouaux, 1998). Hence, our study of the inter-protomer interfacial interactions which may either stabilize or destabilize the large *α*-PFT ClyA pore may yield insights into general factors that could stabilize higher order *α*-helical assemblies. In particular, unlike the AHL pore, we discovered that a set of persistently interacting residues, which were buried in the inter-protomer interface, showed destabilizing energetics, mostly because of desolvated charges that were buried in the low-dielectric protein interior (Figure 6*a*). Selective mutagenesis of these destabilizing residues could be performed to engineer the interfacial stability of ClyA while retaining the gross structural conformation of the pore as well as protomer-protomer conformational specificity, similar to the ClyA mutants produced by directed evolution (Soskine et al., 2013). Stable, rationally engineered mutant nanopores could then be employed for more robust nanopore technologies.

Our algorithm has limitations. Classical MD force-fields with fixed charges, such as the Amber variant employed in this study, do not include polarizability and therefore can frequently over-stabilize salt bridges (Debiec et al., 2014). Thus, while the membrane certainly influences pore conformations, it is not trivial to tease apart the relative contributions of the force-field versus the membrane-environment towards an increased number of salt bridges and/or occurrence frequencies observed in the pore simulations. Secondly, while calculating per-residue non-bonded energies for screening criteria 3, we only compute the enthalpy and neglect entropic contributions. Therefore, our energetic estimates are not free-energies. We deliberately chose to this route because the entropic contributions of residues were intrinsically linked to solvent energetics, and thus would have required computationally expensive enhanced sampling simulations to estimate. This would have negated some of the main advantages of our algorithm – speed and automatability. Enthalpy contributions, which were easy to assess, were arguably of sufficient accuracy to serve as screening criteria. For example, in the directed evolution experiments of Soskine *et al.* to produce stable mutant ClyA pores (Soskine et al., 2013), one of the stabilizing mutations was swapping of the charged E103 to G. In our analysis of per-residue non-bonded energies, E103 was highly destabilizing with an interfacial energy of +78.1 *±* 11.8 kcal/mol, consistent with the experiments of Soskine *et al.* (Soskine et al., 2013). Thirdly, the algorithm predicted residues that were important for stabilizing inter-protomer interface in the pore conformation. The algorithm cannot estimate the effects or even the occurrence of these residue interactions in other conformations of PFTs, whether water-soluble monomer, membrane-bound oligomers or pre-pores. We have previously shown through extensive molecular simulations that the inter-protomer interfaces of membrane-inserted ClyA pore intermediates are similar to that of the fully formed pore (Desikan, Maiti, & Ayappa, 2017). However, for *β*-PFTs such as AHL, conformations of the pore intermediates, pre-pore and the fully formed pore are known to be different (Peraro & van der Goot, 2016; Los et al., 2013). Despite these limitations, the fact that the algorithm was able to predict some of the critical, experimentally validated, inter-protomer hydrogen bonds and salt bridges in both ClyA and AHL suggests that if not all, at least a subset of important protomer-protomer interactions predicted by our algorithm may be common across conformations. These interactions could then serve as potential targets towards inhibiting PFT action.

In conclusion, we successfully employed and validated an *in silico* algorithm for accurate identification of inter-protomer stabilizing electrostatic interactions in membrane-inserted PFT pores. Apart from predicted residues whose mutations were already known to be detrimental towards pore stability, the algorithm also predicted a novel set of residues in both cytolysin A and *α*-hemolysin pores, that could potentially serve as targets for inhibitors which could halt pore formation. From a computational perspective, since the algorithm employs classical equilibrium MD simulations and serially applied, knowledge-based criteria, we recognize that our algorithm could easily be automated and integrated into a parallel high-throughput pipeline for efficient computation.

## 4. Materials and Methods

### 4.1. Fully atomistic equilibrium MD simulations of the ClyA and AHL pores in DMPC membranes

Detailed system set-up and all-atom simulation protocols for both ClyA and AHL pores in DMPC lipid membranes have been previously described (Desikan, Patra, et al., 2017). Briefly, both pores were inserted into pre-equilibrated 18 nm*×*18 nm DMPC membranes, solvated with TIP3P water (Jorgensen et al., 1983), Na^+^ and Cl^−^ ions to a physiologically relevant 0.15 M NaCl, and subjected to 500 ns long unrestrained production runs in the NPT ensemble at 310 K (stochastic rescaling thermostat (Bussi et al., 2007)) and 1 bar (semi-isotropic Parrinello Rahman barostat (Parrinello & Rahman, 1981)). Particle mesh Ewald electrostatics (real space cut-off of 1.0 nm) (Darden et al., 1993) and the leap-frog integrator with a 2 fs integration time step were employed, and all bonds were constrained using the LINCS algorithm (Hess et al., 1997). The AMBER ff99SB-ILDN-*ϕ* force-field (Nerenberg & Head-Gordon, 2011; Lindorff-Larsen et al., 2010) was used for the protein atoms and the ‘Slipid’ parameters were used for DMPC (Jämbeck & Lyubartsev, 2012). MD simulations were performed using GROMACS v*4.6.4* (Pronk et al., 2013), images were rendered with VMD *1.9.4* (Humphrey et al., 1996), and graphs were plotted with MATLAB *R2018a*.

### 4.2. Analysis

Hydrogen bond and salt bridge analysis was carried out using the timeline tool in VMD. Crystal structure salt bridges and hydrogen bonds (Figure 3*c, d*) were computed from the PDB structures of ClyA and AHL after addition of hydrogens according to the force-field by using the pdb2gmx tool in GROMACS. Per-residue solvent accessible surface areas (SASA) of residues were computed using the GROMACS SASA analysis tool (Eisenhaber et al., 1995). Buried and exposed residues were identified by computing the relative solvent accessibility (RSA (Tien et al., 2013) of the residues from SASA; a residue was considered to be buried if its RSA was *≤* 25%. (Levy, 2010) The occurrence frequency of a salt bridge (shown in Figures 4 and 5 for ClyA and AHL) were defined as *N*_*on*_*/N*_*total*_, where *N*_*on*_ is the number of frames in which the salt bridge is formed, and *N*_*total*_ is the total number of frames in the ensemble. Per-residue non-bonded interaction energies of a residue in the inter-protomer interface, Δ*E*^*vdW*^ + Δ*E*^*Coul*^, were computed by calculating the interfacial energy between two protomers using the g mmpbsa tool (Kumari et al., 2014; Baker et al., 2001; Pronk et al., 2013) and then decomposing it into residue-wise contributions. For analysing per-residue energetics as shown in Figure 6*a, b, N*_*total*_ = 1500 frames from 470 *−* 500 ns of the ClyA and AHL MD trajectories were analysed with g mmpbsa, and the protein dielectric constant was assumed to be *ϵ*_*p*_ = 2. Error bars reported in Figure 6*a, b* are standard errors of the mean of the *i*^*th*^ residue computed across all *N* interfaces of the homo-oligomeric ClyA and AHL pores. For estimating the amino acid frequencies across homologous protein sequences of ClyA and AHL, we used the ConSurf web-server (Landau et al., 2005). We obtained aligned homologous protein sequences using the HMMER search algorithm on the UniProt database, and then carrying out multiple sequence alignments with the MAFFT-L-INS-I algorithm. 7 and 10 aligned homologous sequences were obtained for ClyA and AHL, respectively, and amino acid occurrences were depicted as protein sequence logos using the WebLogo server (Crooks, Hon, Chandonia, & Brenner, 2004).

## Acknowledgement(s)

The authors thank Sandhya S. Visweswariah and Raghavan Varadarajan for discussions.

## ABBREVIATIONS

AHL: *α*-hemolysin
ClyA: Cytolysin A
DMPC: 1,2-dimyristoyl-sn-glycero-3-phosphocholine
PDB: Protein data bank
PFT(s): Pore-forming toxin(s)
MD: Molecular dynamics
RSA: Relative solvent accessibility
RMSD: Root mean square deviation
SASA: solvent accessible surface area

